# The mechanical code of DNA impacts its interaction with DNA gyrase

**DOI:** 10.1101/2025.10.21.683783

**Authors:** Bailey G. Forbes, Rosella Pinckney, Sisi Liu, Aditi Biswas, Heather Potter, Elizabeth R. Morris, Aakash Basu-Biswas

## Abstract

DNA:protein interactions involving large structural deformations of DNA underpin essential biological processes. Although correlative evidence suggests that local, sequence-encoded, mechanical properties of DNA can modulate its interactions with large bent-DNA complexes like nucleosomes, direct high-throughput measurements of programmable mechanical modulation of DNA:protein interactions remain lacking. DNA gyrase is a type II topoisomerase that introduces negative supercoils in bacterial chromosomes *via* a process that involves extensive, nucleosome-scale, DNA wrapping around its two C-terminal domains (CTDs). Here we combine Systematic Evolution of Ligands by EXponential enrichment (SELEX), neural network predictions of DNA intrinsic cyclizability, and high-throughput DNA-compete binding assays to broadly reveal that sequence-encoded DNA mechanics tunes gyrase:DNA interactions by modulating wrapping of DNA around the CTDs. Further, we find that both genomic gyrase cleavage sites, and SELEX-enriched strong gyrase-binding sequences, display marked mechanical asymmetry: an extended region of flexible DNA facilitating wrapping around only one CTD exists on one half of the enzyme footprint. High throughput binding assays further reveal that strong binding to one CTD alone can compensate for weaker dual binding, suggesting that asymmetric attachment may have evolved to balance the need for stable anchoring with conformational flexibility required for catalytic remodelling. Additionally, we identify key GC-rich motifs that independently enhance gyrase:DNA interactions, also in an asymmetric fashion. Our findings establish sequence-encoded DNA mechanics as a tunable determinant of protein:DNA interactions and illustrate how functional asymmetry within a symmetric enzyme can couple stable substrate association with structural plasticity.

## INTRODUCTION

### DNA mechanics impacts protein:DNA interactions

Local mechanical distortions of DNA, such a bending, wrapping, melting, or kinking, ubiquitously accompany protein:DNA interactions^1^. Examples include the wrapping of DNA around nucleosomes^2^, tight loop-formation by transcription factors^3^, or DNA bending by DNA-repair factors and topoisomerases. Consequently, the local mechanical pliability of DNA to accommodate such mechanical distortions might impact critical DNA:protein interactions and their downstream processes^1^. Indeed DNA-mechanics has been shown to pre-pay the energetic penalty associated with the binding of diverse transcription factors and repair factors, and DNA sequences that are intrinsically rigid or flexible have been shown to impact nucleosome formation^4^. Further, computational^5–7^, single-molecule^8,9^, and high-throughput studies^4,10,11^ have all suggested that local DNA structure and mechanics are sequence-dependent, leading to the hypothesis that selection for DNA mechanics via a “mechanical code”^10^ may facilitate or hinder specific DNA:protein interactions at various genomic *loci*.

A challenge for testing this hypothesis has been the historic lack of high-throughput methods to measure the sequence-dependence of local DNA mechanics, i.e. to decipher the mechanical code. We recently introduced intrinsic cyclizability – a measure of mesoscale DNA mechanics related to bendability^10,11^ – which captures how rapidly a short 50 bp segment of DNA can cyclize via bending or other deformations. We developed loop-seq (Fig. 1a) – a high throughout assay to measure the intrinsic cyclizabilities of up to ∼100,000 pre-specified 50 bp DNA sequences at a time that can be chosen at will to, for example, tile a long region of the genome. Because of the loop-seq assay, intrinsic cyclizability remains the only mesoscale mechanical property of DNA whose sequence-dependence can be measured in high-throughput. Further, we trained neural networks based machine learning models on the basis of loop-seq measurements to be able to predict intrinsic cyclizability from any input 50 bp DNA sequence^10,12^. Our studies revealed how sequence-encoded DNA mechanics has impacted nucleosome organization – which involves extensive mechanical distortions of DNA – around various genomic *loci* in diverse organisms^10–12^.

**Figure 1:**
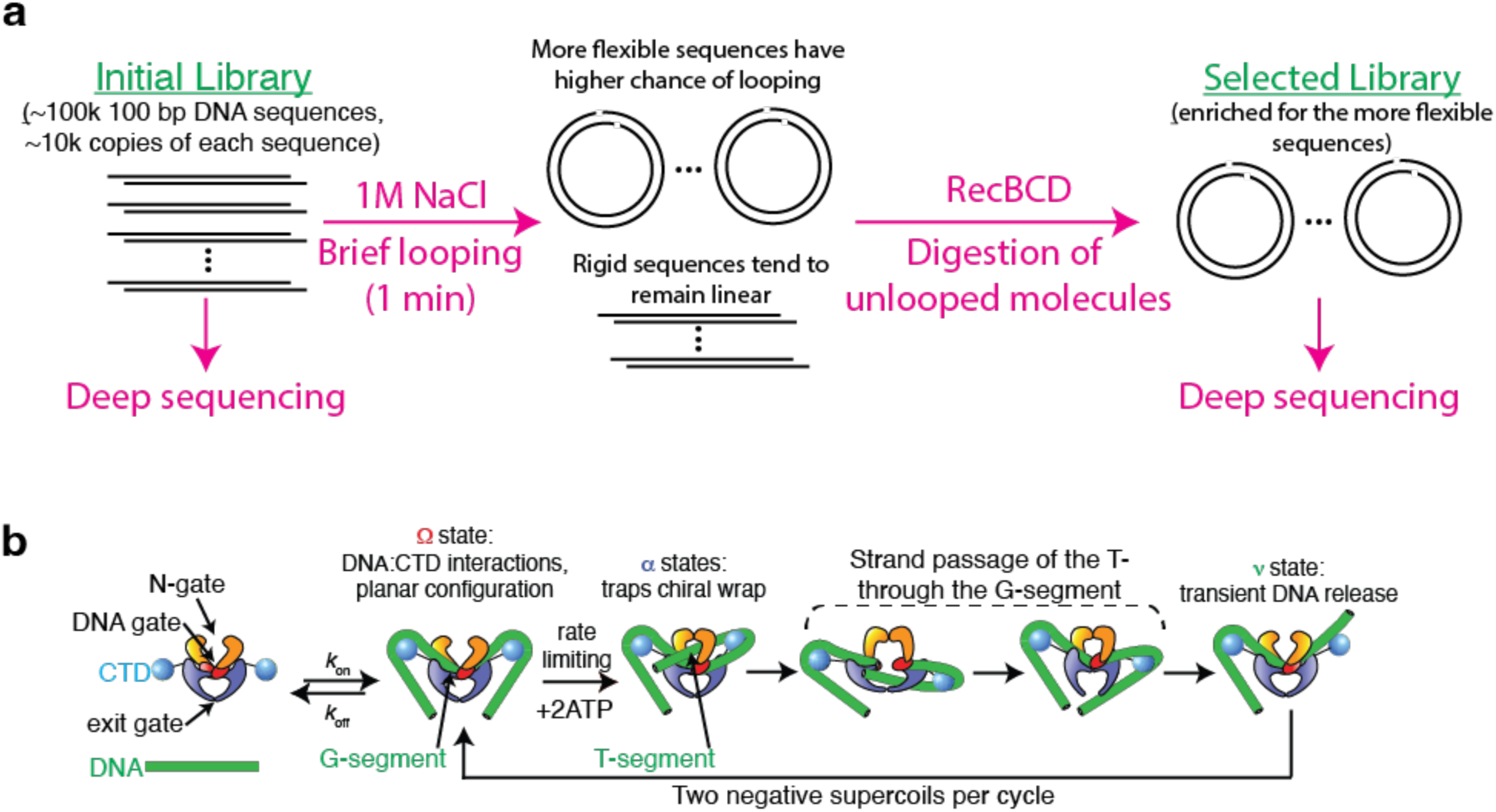
(a) Schematic of the loop-seq protocol^46^. An initial library containing several thousand copies each of ∼100,000 pre-specified, short (50 bp), DNA sequences is chemically synthesized. Molecules are processed to have stickly single-stranded overhangs. The library is permitted to undergo brief intramolecular cyclization via annealing of the ends and uncyclized molecules are enzymatically digested. The original library and the selected library enriched for the more cyclizable molecules are deep-sequenced. For each sequence, the logarithm of the ratio of its relative population in the selected pool to that in the original pool is a measure of cyclizability. **(b)** Schematic of the structural intermediates of the gyrase:DNA complex visited during processive supercoiling, based on our earlier single-molecule works^21,22^ and related works. The gyrase holoenzyme contains three protein:protein interfaces, or gates: the DNA gate which binds and cleaves duplex DNA, the N-gate formed by the ATPase domains, and an exit gate. Gyrase loads on to DNA in the Ω state, with DNA extensively wrapped in a planar fashion around the CTDs and a segment of DNA (G-segment) bound to the DNA-gate. Ω remodels to the α state via a rate-limiting rearrangement of the trapped DNA contour into a chiral loop. A section of the loop (the T-segment) now lies within the N-gate. Cleavage of the G-segment via opening of the DNA gate and the passage of the T-segment through it are accelerated by ATP hydrolysis. The passed T-segment is expelled through the exit-gate, and DNA is transiently released from one or both CTDs (the ϖ state) before the enzyme resets to Ω.

### DNA gyrase wraps DNA on the scale of the nucleosome

DNA gyrase is a bacterial protein that wraps and bends DNA on a scale similar to that of the nucleosome. It is a heterotetrametric bacterial type II topoisomerase of the A_2_B_2_ form, and the only known enzyme to negatively supercoil DNA^13,14^. Negative supercoiling of DNA by gyrase is central to bacterial physiology: it compacts the genome via plectoneme formation, relaxes supercoils ahead of transcription and replication forks, delinks daughter chromosomes, facilitates processes that require access to single-stranded DNA such as transcription and replication initiation, and serves as a secondary messenger communicating environmental conditions to gene expression programmes. Because of its critical role in all bacterial life, gyrase is an important antibiotic drug-target.

Directional supercoiling is achieved through chiral wrapping of DNA around the C-terminal domain (CTD) of the A subunit of gyrase (GyrA), forming a DNA loop of positive chirality in the path of DNA. Gyrase subsequently cleaves the DNA, creating a double-strand break to pass the loop through itself. This inverts the chirality of the loop and leads to the introduction of two negative supercoils. Bulk experiments clearly show that isolated CTDs can trap positive loops along DNA^15,16^, and smFRET experiments have revealed that the CTDs are capable of wrapping ∼140 bp of DNA by 180° *in vitro*^17^. Further, hydroxyl radical footprinting experiments reveal a characteristic pattern of protection with a 10 bp periodicity, indicating ∼80 bp of DNA wrapped on the surface of the two CTDs^18^. Thus, the central role of the CTD in chirally wrapping DNA in the shape of a positive loop has long been appreciated. More recently, cryo-EM has enabled direct visualization of the gyrase holoenzyme in complex with DNA, showing expensive wrapping of DNA around the two CTDs^19,20^.

From our single-molecule studies^21,22^ that built on decades of earlier work^14^, we constructed a cycle of structural intermediated visited by the gyrase:DNA complex during every catalytic cycle. These studies revealed a new, on-pathway configuration of the holoenzyme called Ω, in which the CTDs engage DNA in the shape of a planar bend, but does not chirally wrap it in the shape of a loop (Fig. 1b). Further, our single molecule studies show that Ω is the enzyme’s longest-lived state at all [ATP], in turn revealing that extensive DNA:CTD interactions are not restricted to a transient chiral wrapping intermediate. There are, therefore, extensive structural parallels between gyrase:DNA interactions and the nucleosome – the size of the footprint (∼150 bp in either case) and the long-lived bending deformations enforced on DNA via tight wrapping on the surface of the CTD or the histone octamer^2^. Our earlier works involving loop-seq^10,11^, and several other independent works^23^, have long established a functional role of sequence-encoded DNA flexibility in regulating nucleosome organization. Therefore, we hypothesized a similar role in the context of gyrase:DNA interactions.

This hypothesis is also motivated by other observations. The MuSGS – a Strong Gyrase Site found at the centre of the Mu phage genome, has long been speculated to aid gyrase supercoiling and processivity because of sequence features like periodic GC content oscillations that might make the DNA locally flexible, and therefore amenable to wrapping by the CTD^24,25^. Periodic GC patterns have been shown in SELEX-based cyclization assays to be a feature of cyclizable DNA molecules^26^, and are also reminiscent of similar patterns that exist around dyads of nucleosomes where extensive DNA bending (around the histone octamer core) occures^27^. More recently, high-throughput mapping of the cleavage sites of gyrase along the *E. coli* genome in presence of various antibiotics also found a distinct periodic pattern of GC content on either side of cleavage sites^25^, suggesting a role of DNA bendability around cleavage sites, likely assisting with CTD-mediated wrapping of DNA.

A key deficiency, however, in determining whether DNA sequence, *via* the mechanical code, can regulate gyrase:DNA interactions, is the lack of high-throughput measurements of gyrase:DNA interaction affinities across a broad range of sequences. Here we address this by performing (1)

Systematic Evolution of Ligands by EXponential enrichment (SELEX) to identify sequences from a large random DNA pool that strongly bind gyrase and characterize their mechanical properties, and (2) DNA-compete assays to directly measure the gyrase binding propensities of a large number of pre-specified short DNA sequences. Using our predictive neural-nets based model for the sequence-dependence of intrinsic cyclizability, we reveal that sequence-encoded DNA mechanics plays a central role in governing gyrase–DNA recognition and stability, providing direct evidence for the mechanical code as a regulator of topoisomerase function.

## METHODS

### Protein expression

GyrA and GyrB proteins were produced and the GyrA-GyrB complex assembled according to a protocol adapted from Vanden Broeck et al 2019^19^ and Michaelczyk et al 2023^28^.

Plasmids pET28b-EcGyrATWS and pET28b-EcGyrBTWS, produced by V. Lamour (IGBMC and Strasbourg University Hospitals) and gifted by J. Heddle (Durham University), comprised a modified pET28b backbone with the coding sequences for *E. coli* GyrA (2-875) or GyrB (2-875), respectively, inserted between an N-terminal His_10_-tag and a C-terminal Twin-Strep-tag. GyrA and GyrB proteins were expressed in the T1 phage-resistant *E. coli* BL21 (DE3) derivative strain ER2566 grown in Terrific Broth at 37 °C with shaking. Protein expression was induced by addition of 0.5 mM IPTG to log phase cultures (A600 = 0.6) and the cells incubated for a further 18 h at 18 °C. Cells were harvested by centrifugation and resuspended in 50 mL lysis buffer (20 mM HEPES-NaOH pH 7.5, 300 mM NaCl, 1 mM MgCl_2_, 10% (v/v) glycerol, 20 mM imidazole, 0.5 mM TCEP, 1:10,000 Benzonase, and 1× EDTA-free mini complete protease inhibitors (Roche) per 10 g of cell pellet).

### Protein purification

The cell suspension was lysed by sonication and clarified by centrifugation for 1 h at 50,000×g and 4 °C. The supernatant was subsequently applied to a 5 mL HisTrap FF (Cytiva) column on an AKTA Pure 25 M (Cytiva) held at 2-4 °C in a refrigerated cabinet. The column was washed with 10 column volumes (CVs) HisTrap Wash Buffer (20 mM HEPES-NaOH pH 7.5, 300 mM NaCl, 10% (v/v) glycerol, 20 mM imidazole, 0.5 mM TCEP) and step elution applied with HisTrap Elution Buffer (20 mM HEPES-NaOH pH 7.5, 300 mM NaCl, 10% (v/v) glycerol, 375 mM imidazole).

Peak fractions were pooled and loaded at a slow flow rate onto a 20 mL column containing StrepTactin (IBA) resin, followed by washing with 15 CVs StrepTactin wash buffer (20 mM HEPES-NaOH pH 8, 60 mM NaCl, 10% (v/v) glycerol, 1 mM EDTA-NaOH pH 8, 0.5 mM TCEP) and a step elution with StrepTactin Elution Buffer (20 mM HEPES-NaOH pH 8, 60 mM NaCl, 10% (v/v) glycerol, 1 mM EDTA, 2.5 mM desthiobiotin, 0.5 mM TCEP).

Peak fractions were pooled and the His_10_-and Twin-Strep-tags were cleaved overnight at 4 °C using in-house produced His_6_-TEV and GST-3C proteases in a 1:100 protease:protein ratio. Protein was loaded onto two 5 mL HiTrapQ HP (Cytiva) anion exchange columns in series, washed with 20 CVs HGED buffer (50 mM Na-HEPES pH 8, 10% glycerol (v/v), 1 mM EDTA, 0.5 mM TCEP) and a series of increasing 0.1 M NaCl steps were used to elute protein. Peak protein eluted at 0.2 M NaCl was pooled and concentrated to 3-10 mg/mL, snap frozen and stored at-80 °C.

### GyrA-GyrB complex assembly

GyrA was diluted to 2 μM and GyrB was diluted to 10 μM in Storage Buffer (50 mM Tris-HCl pH 7.5, 100 mM potassium glutamate, 10% (v/v) glycerol, 2 mM DTT) and the two were combined in 1:1 volumetric ratio to achieve a 1:5 stoichiometric ratio of GyrA:GyrB. After incubation for 1 h at 4 °C, the 1 μM GyrA-GyrB complex was snap frozen in liquid nitrogen and stored at-80 °C.

## SELEX

We used SELEX to select, over 7 rounds of selection and amplification, the best sequences for gyrase binding from a large pool of approximately 10¹³ unique 167 bp random double-stranded DNA fragments.

The initial dsDNA random pool was prepared from purchased ssDNA. ssDNA with a central random 133 nt region, flanked by constant 17 nt adapters (left adapter: 5’-ATTGCCGTCCGTACCGT-3’; right adapter: 5’-TTGGGTGCGCGAAACGA-3’) were chemically synthesized and PAGE purified (IDT). A single cycle of PCR-based strand elongation with Phusion polymerase (NEB) in presence of just the reverse primer was carried out to produce a dsDNA library, which served as the initial library for SELEX rounds.

For each SELEX round, gyrase tetramer (in tetramer storage buffer) was incubated with the library (amount per round of SELEX (1-7): 35, 17, 17, 10, 10, 4, 4 pmol) at a DNA:tetramer ration of 10:1 for rounds 1-5, and 5:1 for subsequent rounds, in a total reaction volume of 100 µl, in presence of 1x gyrase binding buffer (50 mM tris-Cl, pH 7.5, 55 mM KCl, 10 mM MgCl_2_,5 mM DTT, 5% glycerol), for 1 hour at room temperature. Excess DNA forces the DNA sequences to compete among each other for enzyme binding. Electrophoretic Mobility Shift Assays (EMSA) was performed to separate bound and unbound fractions using a 6% TBE gel in 1X TAE buffer supplemented with 10 mM MgCl_2_ for 2 h at 45 V (Fig. 2b). Gels were stained with SYBR gold, the band corresponding to the bound fraction was cut, and DNA extracted from it using crush and soak. Extracted DNA was PCR amplified using KAPA polymerase (Roche) over 12 cycles. A certain amount of PCR amplified DNA was saved and prepared for Illumina Sequencing, using the Nextera XT index kit, following library preparation sequencing methods for 16S metagenomic sequencing (Illumina). The rest of the DNA was used for the next round of SELEX.

**Figure 2:**
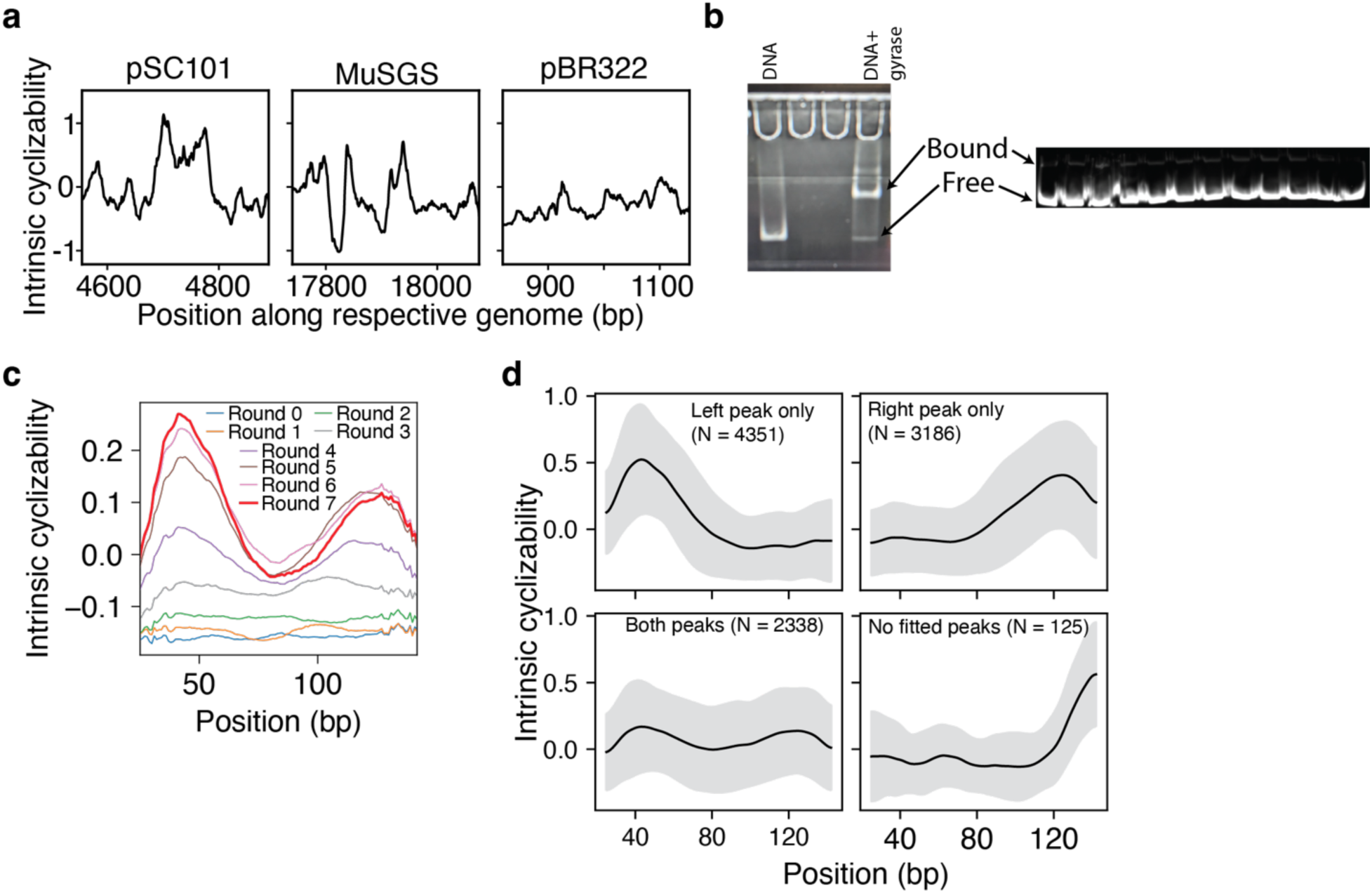
(a) Predicted intrinsic cyclizability as a function of position around known gyrase binding sites in the plasmids pSC101 and pBR322, and in the genome of the bacteriophage Mu. See supplementary note 2a. **(b)** Visualization of bound and free DNA after gyrase binding to a single DNA sequence at 1:1 DNA:gyrase molar ratio (left panel), and to the random library used for SELEX at 10:1 DNA:gyrase molar ratio. **(c)** Mean intrinsic cyclizability as a function of position along the 167 bp DNA fragment, averaged over 10,000 randomly selected unique reads in every round of SELEX. Round 0 refers to sequencing of the original pool of random DNA before any gyrase binding. See supplementary note 2c for plotting details. **(d)** Mean intrinsic cyclizability (black line) ± 1 S.D. (grey band) as a function of position along the 167 bp DNA fragment, averaged over subsets of the 10,000 randomly chosen unique reads from round 7 of SELEX, which were classified as having intrinsic cyclizability peaks on the left side, right side, both sides, and neither sides of the centres of the DNA fragments. See supplementary note 2d for classification and plotting details.

### DNA-compete

DNA compete experiments started with a chemically synthesized library of 1,253 167 bp DNA sequences (Twist Bioscience), which was treated similar to 1 round of SELEX. Gyrase was incubated with the DNA library under low enzyme concentrations, allowing the different DNA sequences to compete for binding. Bound and unbound DNA fractions were separated by gel electrophoresis, and both pools were deep-sequenced. We define binding score of a sequence as the log of the ratio of its counts in the bound pool to that in the unbound pool.

### Sequencing data analysis

Bi-directional sequencing was carried out on an Illumina platform. Reads were merged using fastp. For each round of SELEX, merged reads were first filtered for only those reads that contained the left adapter and the right adapter, and exactly 133 nt between them. Following this, unique reads were identified from each round of SELEX and the counts per mission (CPM) of each unique read was calculated. Reads were sorted according to decreasing CPM. For every round, 10,000 random reads were selected at random for downstream analysis.

## RESULTS

### An asymmetric structural motif for gyrase binding identified *via* SELEX

We used our neural networks based predictive model^12^ for the sequence-dependence of DNA cyclizability to initially establish a potential role of sequence-encoded DNA mechanics in facilitating the wrapping of DNA around the CTDs. We compared predicted intrinsic cyclizability of DNA around three gyrase sites reported earlier in the literature^29^: the Mu phage Strong Gyrase Site (MuSGS), and gyrase binding regions within the plasmids pSC101 and pBR322. pSC101 is known the harbour the strongest gyrase binding site^29^ (Fig. 2a). Consistent with this, the predicted intrinsic cyclizability profile for *pSC101* exhibits two pronounced peaks flanking the center of the cleavage region—features that likely facilitate DNA wrapping around the two CTDs flanking the central cleavage site. The MuSGS site shows a similar but less prominent double-peak pattern, whereas pBR322 lacks discernible peaks. This gradation in prominent flexibility peaks mirrors the known pattern of gyrase binding efficiency to these three sites^29^ (pSC101> MuSGS >> pBR322), and suggests a putative role of sequence-encoded DNA mechanics in regulating gyrase:DNA interactions.

To experimentally identify DNA sequences with high gyrase-binding affinity, we performed Systematic Evolution of Ligands by EXponential enrichment (SELEX) over seven rounds. A starting library comprised approximately 10¹³ unique 133 bp random double-stranded DNA fragments, each flanked by 17 bp constant adapter regions for PCR amplification. Although this represents only a small subset of the theoretical sequence space for 133 bp DNA, it provides extensive diversity for selection. In each SELEX round, gyrase tetramers were incubated with the DNA library under conditions of limiting enzyme, allowing for the selection of the best bending DNA sequences. DNA bound and unbound to gyrase were separated *via* Electrophoretic Mobility Shift Assay (EMSA) (Fig. 2b). Bound DNA was extracted; a small aliquot was sequenced, and the remainder was PCR-amplified to generate input for the next round of selection (7 rounds in total). This iterative process enriched sequences with progressively higher gyrase-binding propensity. From each SELEX round, we randomly sampled 10,000 sequences and used our neural-network model to predict the position-dependent intrinsic cyclizability (DNA flexibility) along each 167 bp DNA fragment. This allowed us to monitor the evolution of DNA mechanical signatures associated with gyrase binding across successive selection rounds.

Analysis of predicted flexibility profiles across SELEX rounds revealed the emergence of a distinct mechanical signature for gyrase binding (Fig. 2c). As selection progressed, selected DNA, on average, had two pronounced peaks in intrinsic cyclizability—one on either side of the central region. These peaks correspond closely to the positions where the C-terminal domains (CTDs) of gyrase are expected to engage the DNA. Our findings suggest that sequence-encoded DNA mechanics is a key determinant of gyrase:DNA interactions, because of the requirement for the gyrase CTDs to extensively wrap DNA. This parallels the role of sequence-encoded flexible DNA at the dyads of most known nucleosomes in yeast in aiding extensive DNA wrapping around the histone core.

Our data reveals an expanded role for DNA wrapping around the CTDs: it contributes to stable binding of gyrase. If stable binding of gyrase to DNA were mediated only *via* G-segment binding to the central DNA-gate (Fig. 1b), it is unlikely that extensive DNA flexibility features on either side of the central G-segment would have been enriched. The observed enrichment in a SELEX experiment that specifically selects for gyrase binding only, therefore suggests that flanking DNA flexibility facilitates enzyme-DNA complex formation via likely engaging the CTDs. As our single-molecule measurements identify Ω (Fig. 1b) as an on-pathway intermediate with extensive DNA:CTD interactions in which the enzyme dwells the longest at all [ATP], it is likely that sequence features allowing stable binding of DNA to the CTD stabilize the Ω state. Stable formation of the long-lived Ω, therefore, likely serves the function of properly anchoring gyrase to DNA.

Further analysis revealed that the enriched mechanical motif for gyrase binding in our SELEX experiments is frequently asymmetric. When analyzing the 10,000 randomly chosen sequences from the 7^th^ (final) round of SELEX, we found that 76% of sequences have just one peak in intrinsic cyclizability along its length, either on the ‘left’ or ‘right’ side of the centre (Fig. 2d). Only 23% have both cyclizability peaks, and these peak heights are significantly reduced as compared to the peak height of DNA sequences that have just one peak. Our findings therefore suggest that to effect high-affinity gyrase:DNA binding, stable engagement of DNA with just one CTD can compensate for weaker engagement with both CTDs, and that most selected gyrase binding sequences adopt this asymmetric strategy. There are several possible functional advantages to this arrangement. Processive supercoiling by gyrase requires large-scale conformational rearrangements involving DNA, three dynamics protein:protein interfaces serving as gates for DNA transfer within the enzyme, and both CTDs. Also, we have previously reported that transient release of DNA from one or both CTDs is necessary, and on-pathway during processive supercoiling^21,30^. Anchoring the enzyme *via* stable binding to one CTD, while maintaining flexible or transient interactions with the other, could therefore facilitate such dynamic transitions. Such asymmetric model of engagement, where one CTD:DNA interaction can compensate for the other to effect stable binding, may represent an evolutionarily optimized balance between stability and plasticity within the gyrase–DNA complex.

### Gyrase:DNA interactions are independently impacted by GC-rich sequence motifs

The parameter intrinsic cyclizability, as defined for 50 bp DNA, is ultimately derived exclusively from its sequence. But the mapping between sequence and flexibility is highly degenerate and complex. Therefore, we analyzed our SELEX data to see if, in addition to selection for flexibility, there are simpler sequence motifs being enriched in the selected pool.

We first analyzed the evolution of the average GC content profile along the 133 bp random region per round of SELEX (Fig. 3a). We overall see periodic oscillations in GC content (Fig. 3a), which strengthens with subsequent SELEX rounds as seen from Fourier analysis of the GC content oscillations (Fig. 3b, left panel). While the average GC content across 10,000 sequences in each round of SELEX shows prominent periodic oscillations only on the “left” half of the sequence (Fig. 3a), we conclude this is owing to proper phasing between sequences (see supplementary note 3b). Individual sequences do display 10 bp periodic GC content oscillations on either side of the centre, as confirmed *via* Fourier analysis separately on the two halves (Fig. 3b, right panel). We can speculate that the phased oscillations on the left half are a result of a phase being imposed by the constant adapter sequence on the left half of the molecule. We further find that periodic oscillations in GC content also largely mimic the asymmetry in DNA flexibility: when the sequences are grouped into those with a flexibility peak on one half of the molecule, the periodic GC-content oscillations are restricted to that half (Supplementary Fig. S1). This suggests a connection between GC content oscillations and DNA flexibility. Indeed, periodic oscillations in GC content have been shown earlier in SELEX experiments to favour DNA cyclization^26^, and are also found on either side of mapped *E. coli* gyrase cleavage sites^25^. Further, 3% of known nucleosome formation regions in yeast also have periodic oscillations in GC content^31^.

**Figure 3:**
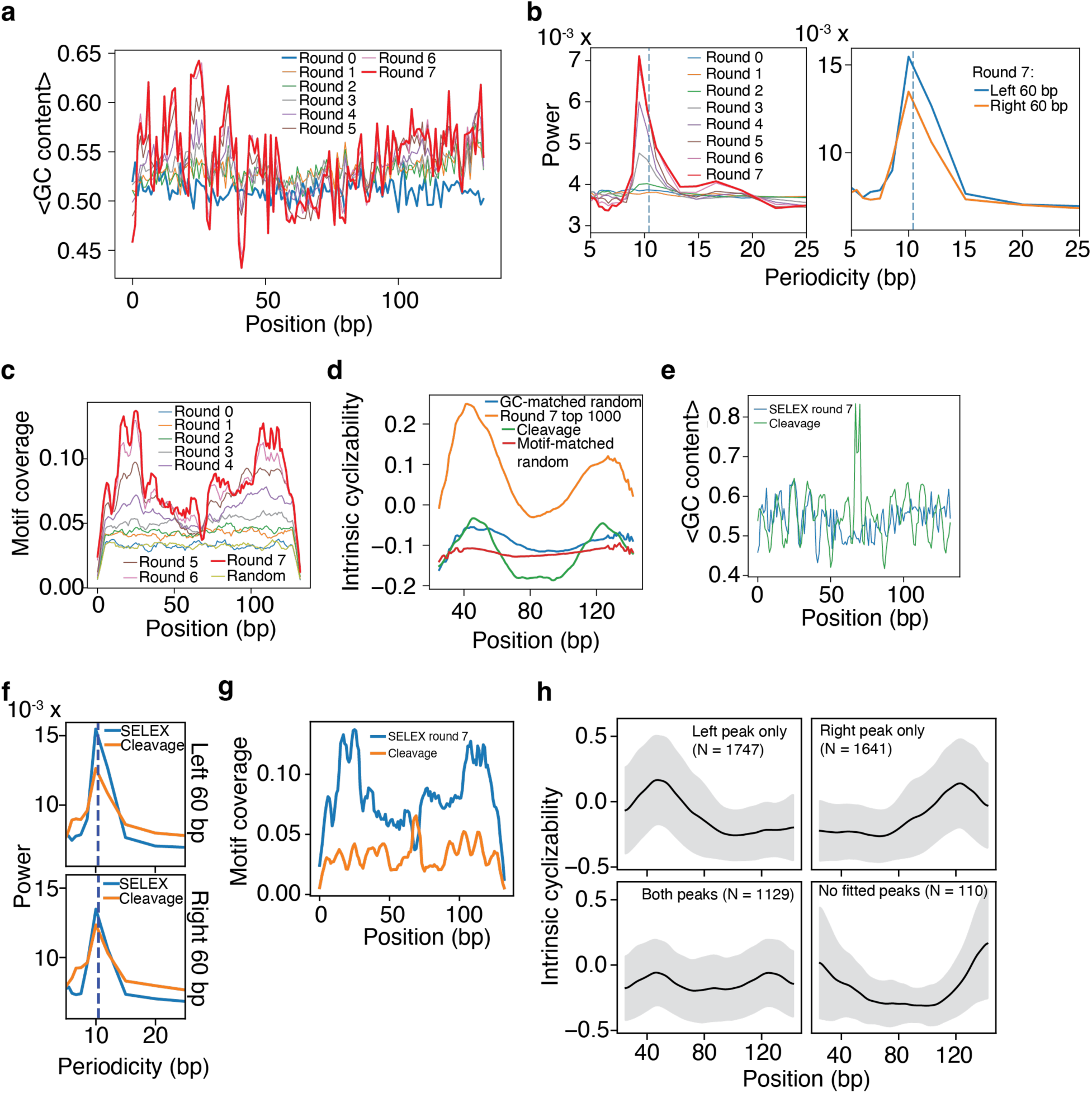
(a) Mean GC content as a function of position, averaged over the 10,000 randomly selected DNA sequences during each round of SELEX. **(b)** Left panel: Fourier power associated with oscillations in GC content along the central 133 bp region, as a function of periodicity, calculated via categorical spectral analysis. For each round, power vs perodcicity was individually calculated for the 10,000 randomly selected sequences, and then averaged. Right panel: Averaged power as a function of periodicity from among the 10,000 sequences in round 7 of SELEX, calculated individually for GC content oscillations within the first 60 bp and last 60 bp of the 133 bp region. See supplementary note 3b for plotting details and categorical spectral analysis. **(c)** Mean motif coverage profile of the 20 identified 6-mers (Table 1) as a function of position along the 133 bp, averaged over the 10,000 randomly selected unique sequences in every round of SELEX. For every round, for each base position, plotted is the fraction of the 10,000 sequences where that base position lies within any occurrence of any of the 20 motifs. **(d)** Mean intrinsic cyclizability of DNA as a function of position along the 167 bp region, averaged over (i) the 10,000 randomly selected sequences from round 7 of SELEX, (ii) a randomized version of these 10,000 sequences that preserve the per-sequence GC profile, (iii) a set of 10,000 otherwise random DNA sequences that have the same, per-sequence, coverage profile of the 20 6-mers as the 10,000 round 7 SELEX sequences, and (iv) the 4,627 identified cleavage sites in E. coli in presence of the antibiotic Cfx. See supplementary note 3d for details. **(e)** Mean GC content as a function of position along the 133 bp variable region of the 10,000 round 7 SELEX sequences, and in the 133 bp region around the mapped gyrase cleavage sites in presence of Cfx. **(f)** Mean fourier power, obtained via categorical spectral analysis of individual sequences followed by averaging, of the 10,000 133 bp variable sequences in round 7 SELEX, and of 4,627 sequences spanning 133 bp around the mapped Cfx-induced gyrase cleavage sites in E. coli. Power was calculated separately for the first 60 bp of each sequence and the last 60 bp of each sequence. **(g)** Average coverage of the 20 identified 6-mer motifs, as a function of position along the central 133 bp region of the 10,000 round 7 SELEX sequences, and in the 133 bp region flanking the 4,627 Cfx-induced cleavage sites in E. coli. **(h)** Mean intrinsic cyclizability (black line) ± 1 S.D. (grey band) as a function of position along the 4,627 133 bp DNA fragments spanning Cfx-induced cleavage sites in E. coli, each sequence being flanked by standard SELEX 17 nt adapters, averaged over subsets which were classified as having intrinsic cyclizability peaks on the left side, right side, both sides, and neither sides of the centres of the DNA fragments. See supplementary note 2d for details.

The above observations raise an important question – is the selection for flexibility profile purely a consequence of a selection for GC content oscillations? For each of the 10,000 sequences randomly chosen from the final round of SELEX, we created a randomized version where individual Gs and Cs are randomly replaced by Gs and Cs (and same for As and Ts). Therefore, we preserve, on a per-sequence bases, the same pattern of GC variations along the sequence. We find that although these generated sequences also show peaks in flexibility at the same locations as the SELEX sequences, the peaks are shallower in magnitude (Fig. 3d). Therefore, SELEX is enriching sequences with higher flexibility peaks than would be expected from just the observed enrichment in GC content oscillations. This confirms that flexibility selection is not a pure consequence of the selection for GC content periodicities.

We then asked the reverse question – will selecting for sequences with specific peaked flexibility patterns automatically result in GC-content oscillations of the type seen in SELEX data? We performed *in-silico* evolution with DNA point mutations to generate 20 random DNA sequences whose flexibility profiles match the mean intrinsic cyclizability profile of the 100 SELEX sequences from round 7 (final round) with the highest CPM and classified to have flexibility peaks only on the left half of the molecule (Supplementary Fig. S2). We find that these 20 sequences also automatically have, on average, a similar oscillatory profile in GC content (Supplementary Figure S2). While it is possible that periodic oscillations in GC content are directly impacting gyrase:DNA interactions, the more likely explanation that explains our data is that GC content oscillations are predominantly a consequence of selection for DNA flexibility patterns, and not the other way around.

We observed that some of the peaks in GC content are flattened (Fig. 3a). Therefore, we investigated whether there are longer GC-rich sequence motifs being enriched in subsequent SELEX rounds. We identified the 20 most enriched 6-mers among 10,000 randomly selected sequences from round 7 compared to a similar set from round 0 (Table 1, Supplementary Note A). These motifs are, overall, GC rich, interspersed with short A/T regions. Plotting the mean coverage density of these motifs along the 133 bp random region, averaged over 10,000 randomly selected sequences from each round, we found that these motifs are predominantly enriched in the same two regions flanking the centres, where the CTDs are expected to engage, and where we see intrinsic cyclizability to be selected for in SELEX (Fig. 3c). As with cyclizability, the coverage density is also asymmetric: sequences classified to have a flexibility peak on one half of the molecule only also have higher coverage density only on that half (Supplementary Fig. S3).

As with GC content oscillations, we asked to what extent selection for GC-rich motifs is linked with selection for specific patterns of DNA cyclizability. We created random DNA sequences that have the same coverage density for these 20 6-mer motifs, on a per-sequence basis, along the 133 bp region, as the 10,000 randomly selected sequences from round 7 of SELEX. We found that these generated sequences do not have any discernible flexibility peaks when compared to the top sequences selected by SELEX (Fig. 3d), suggesting that selection for these motifs is not the cause of selection for flexibility peaks, as was the case with GC content oscillations. However, when we analyzed the DNA sequences that had been generated via *in silico* evolution to match the flexibility pattern of SELEX sequences, we found that the coverage density of these motifs is much lower in the evolved DNA sequences (Supplementary Figure S2). Therefore, selection for flexibility peaks does not automatically result in a selection for these motifs (unlike the case with GC content oscillations). Our data therefore suggests that gyrase:DNA interactions may be independently favoured by elevated intrinsic cyclizability of DNA in the region that interacts with the CTDs, and by the presence of long GC rich regions interspersed by shorter AT regions.

### Best binding sequences do not share the same mechanical properties as DNA around known *E. coli* cleavage sites

We next compared the sequence and mechanics of DNA selected for gyrase binding *via* SELEX, with that of DNA flanking the known mapped cleavage sites in *E. coli* in presence of the antibiotic Cfx. Cfx stabilizes the cleaved state. Firstly, cleavage sites are marked by a specific sequence motif (depending on the antibiotic used), and this is reflected in the plot of mean GC content as a function of position along DNA sequences spanning the cleavage site as a regio of distinctly higher GC content (Fig. 3e). Such a region is absent in the SELEX sequences (Fig. 3e). This indicates that sequence features that best stabilize the cleaved state are not involved, per-se, in efficient gyrase binding, and are therefore not enriched among SELEX sequences.

We found that DNA on either side of mapped cleavage sites in *E. coli* does have two extended peaks in intrinsic cyclizability, reminiscent of SELEX-selected sequences (Fig. 3d). The pattern of GC content also exhibits oscillations around known cleavage sites, with oscillatory power on either side of the cleavage site similar to that in the case of SELEX sequences (Fig. 3f). But the flanking cyclizability peak heights are comparable to what is automatically achieved via randomizing SELEX sequencing while preserving the G/C patterns, i.e., by simply preserving the existing GC content distributions (Fig. 3d). Further, the coverage density of the 20 6-mer motifs identified to have been enriched in SELEX sequences are also less prevalent around gyrase cleavage sites (Fig. 3g). Thus DNA sequences flanking *E. coli* loci that have evolved to be most stably cleaved by gyrase have not necessarily evolved the mechanical patterns to optimise gyrase binding. This may reflect a need for less tight CTD binding in order for gyrase to remodel DNA into a chirally wrapped and eventually cleaved conformation amenable for supercoil introduction. This may also reflect biological function: perhaps *E. coli* has specifically not evolved strong gyrase binding sites so that the ∼300 gyrase molecules within a cell^32^ can more quickly sense topological strain when diffusing, than they can if stably anchored somewhere.

### Gyrase cleavage sites in *E. coli* also have an asymmetric structural and sequence motif

Is the mechanical asymmetry seen in SELEX-enriched sequences also prevalent among known gyrase binding sites in *E. coli*? We found that intrinsic cyclizability patterns are indeed asymmetric for 73% of the previously mapped Cfx-induced cleavage sites in *E. coli*, and symmetric only for 24% (Fig. 3h). Strength of periodicities in GC content on either side of the cleavage sites also mimic the asymmetry in DNA flexibility (Supplementary Figure S4). These results show that asymmetric flexibility patterns on either side of the centre, as seen in SELEX sequences, are also prevalent around actual gyrase cleavage sites in *E. coli*, despite these sites have far lower overall DNA cyclizability. It supports our hypothesis that strong binding of one CTD to DNA might compensate for weaker binding of both, allowing for the evolution of sites with asymmetric mechanical patterns to balance the need for stable gyrase anchoring with internal dynamic plasticity.

### Direct high-throughput quantification of binding propensity of gyrase to various DNA sequences

SELEX progressively enriches a pool of DNA with sequences with high gyrase affinity. It does not, however, provide a quantitative measure of binding strength of any individual DNA sequence to gyrase. Therefore, though claims such as higher flexibility peaks enabling stronger binding, or strong binding of one CTD compensating for weaker binding of both, are suggestive based on the SELEX data, they need to be independently verified through experiments that directly measure the binding strength of individual DNA sequences to gyrase.

To measure binding propensity, we performed high-throughput DNA-compete assays (methods) on a pool of chemically synthesizes 1,253 167 bp DNA sequences. Only the central 133 bp were variable among the different sequences. The flanking 17 bp were constant, as for the SELEX experiments, and used for PCR amplification. These 1,253 sequences included:

1. Top SELEX-derived binders – the hundred most enriched sequences from round 7 of SELEX classified as having a single peak in intrinsic cyclizability on the left side of the molecule, and the 100 top-sequence with two peaks.
2. Synthetically designed cyclizability variants – 400 sequences designed to have specific intrinsic cyclizability variations along their lengths. We first defined 10 intrinsic cyclizability profiles possessing a single peak in intrinsic cyclizability on the left half of the molecule, with peak location and width matching the top 100 SELEX-selected sequences from round 7 classified as having a single peak on the left side (Fig. 4a, solid line). We varied the peak heights across the 10 profiles, with the shallowest peak heights dipping below the baseline intrinsic cyclizability and peak heigh being variable across the 10 profiles. For each profile, we generate, at random, a group of 20 DNA sequences across which the mean intrinsic cyclizability as a function of position closely matches the profile (Fig. 4a). This was done through *in silico* point mutation and selection (see Supplementary Fig. S2). Figure 4a shows the mean flexibility as a function of position, averaged over the 20 sequences of each group. Similarly, 10 additional profiles with two flexibility peaks in each profile were used to generate 10 more groups (groups 11-20) of 20 sequences each, whose mean flexibility as a function of position is shown in Figure 4a.
3. Random DNA sequences – 400 DNA sequences for which the sequence of the central 133 bp region was chosen completely at random.
4. Cleavage site flanking sequences – 250 sequences derived from genomic regions flanking previously mapped gyrase cleavage sites identified in the presence of ciprofloxacin (Cfx)^25^. Among these, the first 10 sequences corresponded to those scoring highest for cleavage propensity, the next 10 scored lowest, the following 10 had the highest read counts (N3E) associated with cleavage, the next 10 had the lowest read counts, and the remaining 210 were randomly selected from cleavage sites not represented in the previous subsets.
5. Reference controls – three specific sequences: *MuSGS*^29^, the ideal consensus cleavage sequence^25^, and a scrambled version of the consensus sequence^25^.

**Figure 4:**
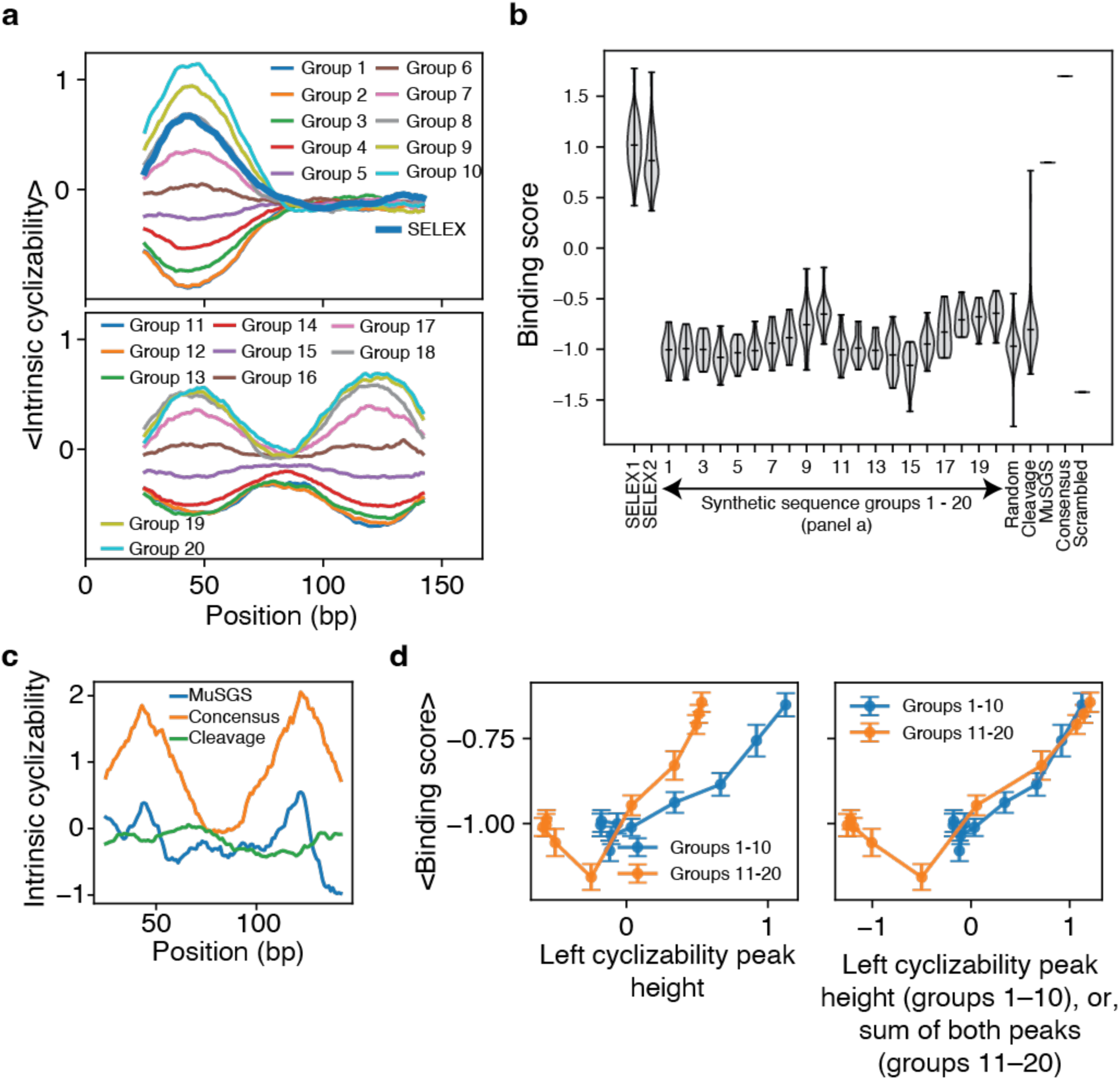
(a) Mean flexibility as a function of position along the 167 bp region, averaged over the 20 synthetic sequences belonging to each group. Thick line: mean intrinsic cyclizability as a function of position, averaged over the 100 sequences from round 7 of SELEX that had the highest counts and were classified as having just a single cyclizability peak on the left side of the molecule. **(b)** Violin plot of DNA binding scores of all categories of DNA sequences. SELEX1 and SELEX2 refer to the 100 most enriched sequences each in round 7 of SELEX that were classified as having a left only intrinsic cycliability peak, and having peaks on both sides respectively. **(c)** Intrinsic cyclizability as a function of position along the 133 bp DNA sequences from the centre of the MuSGS sequence, the previously constructed perfect consensus cleavage sequence^25^, and a scrambled version of the consensus cleavage sequence^25^. Each sequence was flanked by the 17 bp left and right SELEX adapters before intrinsic cyclizability was computed along the sequences. **(d)** Left panel: mean DNA binding score averaged over the 20 sequences in each group, plotted against the mean flexibility in the 10 bp region surrounding the peak location of the 20 sequences within the group. Error bars are s.e.m. Right panel: For groups 1-10, the plot is identical to the left panel. For groups 11-20, the x-coordinate of every point represents the sum of mean flexibilities in the two 10 bp regions surrounding both peak locations of the 20 sequences within that group. See supplementary note 4c for plotting details.

### Binding propensities of different categories of DNA sequences are partly determined by DNA flexibility

DNA binding scores for all sequence categories are summarized in Fig. 4b. As expected, within the two groups of synthetic DNA sequences, increasing the height of the flexibility peak leads to a corresponding increase in gyrase binding score, revealing a direct link between DNA mechanics—encoded by the mechanical code—and gyrase affinity (Fig. 4b). Sequences derived from SELEX show significantly higher binding scores than any other categories of sequences, including the synthetic sequences in groups 1-20 with some of which top SELEX binders share the same intrinsic cyclizability profiles (Fig, 4b). This indicates that intrinsic cyclizability alone does not fully determine gyrase binding, and that other sequence features—such as extended G/C-rich motifs—likely contribute.

Also, DNA fragments flanking known gyrase cleavage sites show binding scores that are, on average, not significantly different from random DNA, although a subset exhibits notably higher scores (Fig. 4b). This is consistent with our earlier observation that intrinsic cyclizability peaks flanking cleavage sites are comparable to what is expected on the basis of GC content oscillations alone, and far shallower than those in SELEX-enriched sequences.

Finally, as expected, the MuSGS has a high binding score. The ideal consensus cleavage sequence, as constructed earlier from the conserved motifs averaged around known cleavage sites^25^, also has a high binding score (despite DNA around cleavage sites on average not being strong binders). We note though that this consensus sequence does not exist in nature, and has very high peaks in cyclizability (Fig 4c). A scrambled consensus cleavage sequence^25^ binds weakly, also consistent with it having non-discernible cyclizability peaks (Fig. 4c).

### Strong DNA binding to one CTD can compensate for weaker binding to two

For the single cyclizability peak series (Groups 1–10), the mean binding score remains approximately constant across Groups 1–5, where the “left” flexibility peak lies below baseline (Fig. 4d, left panel). This plateau likely reflects a constant level of interaction between a gyrase CTD and the right half of the molecule, where flexibility is held fixed across these groups (Fig. 4a). In Groups 6–10, the left-side flexibility peak progressively increases, and so does the binding score, showing that enhanced local flexibility strengthens gyrase binding (Fig. 4d, left panel).

For the double cyclizability peak series (Groups 11–20), binding scores are generally higher than for single-peak sequences with the same left-side peak height (Fig. 4d, left panel). This suggests that the additional flexibility peak on the right half enhances binding. Indeed, this effect appears roughly additive: sequences whose sum of left and right flexibility peak heights equals the single-peak height of Groups 5–10 exhibit comparable binding scores (Fig. 4d, right panel). These observations provide direct support for our hypothesis that strong engagement of one CTD can compensate for weaker engagement of the other, consistent with the asymmetric flexibility profiles observed in SELEX-selected sequences (Fig. 2d), and at mapped gyrase cleavage sites (Fig. 3d).

### Binding preference depends on DNA flexibility and GC content independently

To identify determinants of gyrase binding in a sequence unbiased manner, we analyzed the binding scores of the 400 random DNA sequences in our library. Binding scores were positively correlated with the maximum intrinsic cyclizability along the 167 bp region of each DNA fragment, indicating that indeed mechanical flexibility of DNA, as determined by sequence via the mechanical code – can measurably impact gyrase:DNA binding (Fig. 5a). We also found binding scores to be negatively correlated with overall GC content along the DNA (Fig. 5a), whereas SELEX sequentially enriches for DNA with slightly higher GC content each cycle (Fig 3a). This is possibly because of the independent selection for GC rich motifs, which are largely absent in random DNA. Importantly, GC content and flexibility were themselves uncorrelated (Fig. 5a), indicating that these two sequence properties contribute independently to gyrase binding, consistent with our earlier findings.

**Figure 5:**
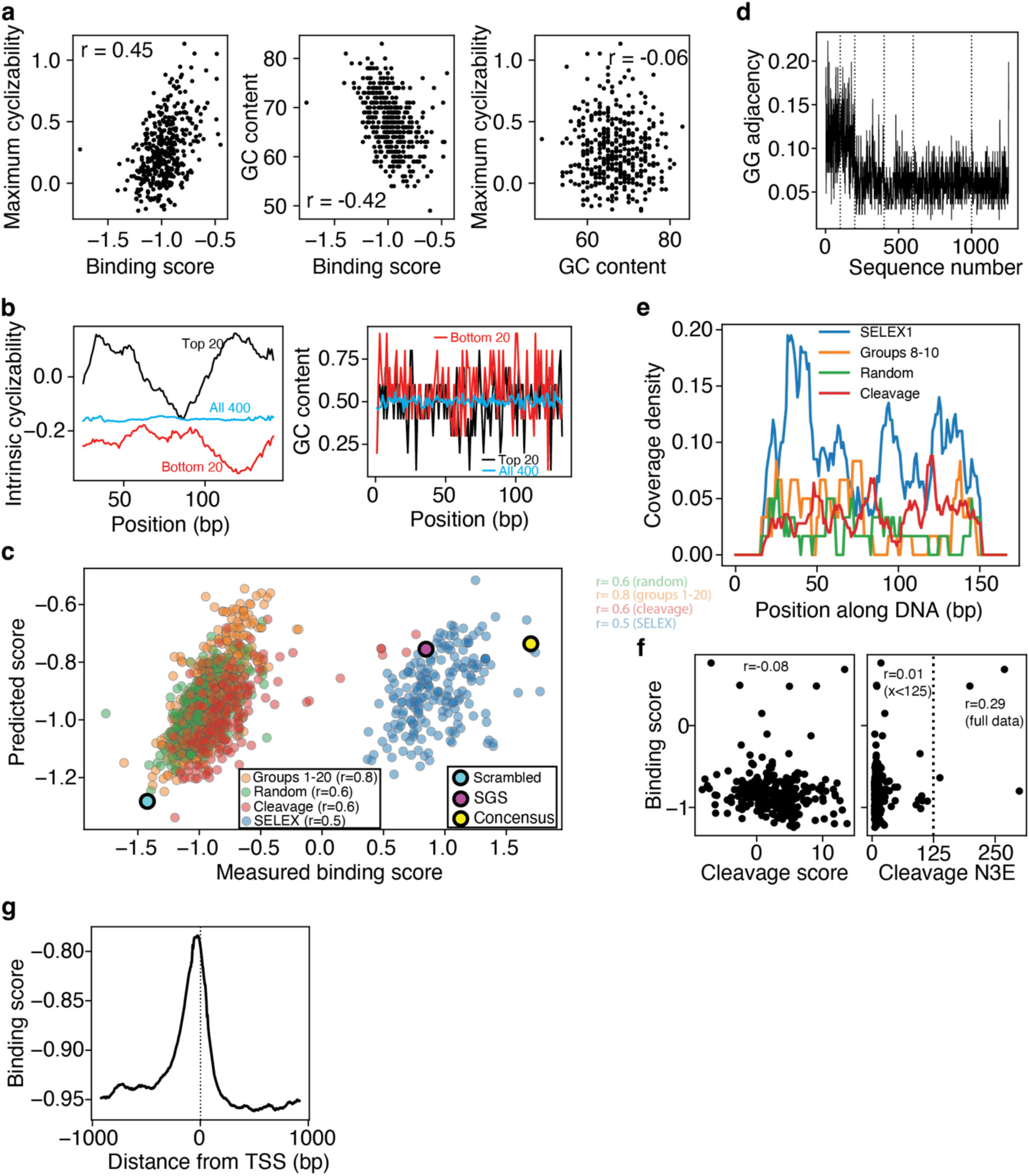
(a) Scatter plots of the 400 intrinsic cyclizability values, binding scores, and overall GC content (number of Gs and Cs in the central 133 bp region) of the 400 random sequences in the library of 1,253 sequences. All possible pairwise scatter among the three variables are plotted. **(b)** Mean intrinsic cyclizability as a function of position (left panel) and mean GC content as a function of position, averaged over the 20 sequences with the highest and lowest binding scores among the 400 random DNA sequences, and also over all 400 sequences. Intrinsic cyclizability was plotted for the full 167 bp region, whereas GC content is plotted for the central 133 bp region, ignoring the constant flanking adapter sequences. Plots are obtained as in Fig. 2c and 3a. **(c)** Predicted (based on equation 1) vs measured binding scores of all the 1,253 sequences, colour coded depending on category. Pearson’s r for the correlation between predicted and measured binding scores for the four categories of DNA sequences is indicated. **(d)** GG adjacency rate per sequence: for every sequence, plotted is the fraction of all dinucleotides within it that are ‘GG’. **(e)** Mean motif coverage profile of the 20 identified 6-mers (Table 1), as a function of position along the central 133 bp region, averaged over the various categories of sequences within the library. In each category, for each base position, plotted is the fraction of the sequences within that category where that base position lies within any occurrence of any of the 20 motifs. **(f)** Scatter plot of the binding score of the 250 sequences in the library that were designed to flank 250 previously characterized Cfx-induced cleavage sites in E. coli, plotted against the cleavage score and the number of 3’ ends (N3E, a measure of cleavage strength) as identified earlier^25^. **(g)** Model prediction of gyrase binding score as a function of distance from Transcription Start Sites, averaged over around 1,797 previously mapped TSSs in E. coli^47^. See supplementary note 5g for plotting details.

When the 400 random sequences were ranked by binding score, the top 20 binders displayed a clear two-peak cyclizability profile centered around the 133 bp region, whereas the bottom 20 were globally less cyclizable (Fig. 5b, left). This is consistent with our findings thus far that DNA cyclizability or flexibility likely aids gyrase binding by allowing easier wrapping around the CTDs. In contrast, GC-content profiles were less distinct between high-and low-binding sequences (Fig. 5b, right).

We next modelled the binding score of each 133 bp sequence as a linear combination of the maximum cyclizability value along the sequence and its overall GC content (see supplementary note B). Applying the fitted model to predict binding scores for all other sequence classes revealed strong agreement between predicted and measured values, with pearson’s r values ranging from 0.5 to 0.8 (Fig. 5c). The only major exception was the set of SELEX-enriched sequences, which consistently exhibited higher binding scores than predicted based on their flexibility and GC content alone. Nonetheless, even within the SELEX set, the relative trend in the data was captured by the model (r = 0.5).

The systematic deviation of SELEX sequences from model predictions suggests that additional sequence-level features enhance their binding beyond what is explained by DNA mechanics or overall GC composition. As noted previously, SELEX-derived binders are specifically enriched in GC-rich motifs separated by short AT-rich linkers—patterns that neither arise solely from mechanical selection nor fully explain the observed flexibility. These motifs, which we found to appear less frequently in other sequence groups (Fig. 5d-e), likely encode specific recognition elements that cooperate with DNA mechanics to promote high-affinity gyrase binding.

### DNA around *E. coli* cleavage sites has not evolved to optimize gyrase binding

When we compared our measured binding scores with independently reported cleavage-site scores and cleavage read counts (N3E) in *E. coli*^25^, focusing on the 250 sequences in our library that flank previously mapped gyrase cleavage sites (identified in the presence of ciprofloxacin, Cfx), we observed no significant correlation between binding score and cleavage read count, except for 4 sequences that had very high binding scores and cleavage read counts (Fig. 5f). Thus, the propensity for DNA to stabilize the cleaved state does not directly mirror gyrase binding affinity. This is also reflected in the fact that DNA sequences spanning known cleavage sites have very similar distributions of binding score to completely random DNA sequences (Fig. 4b). Mapping cleavage events and their relative strengths therefore does not provide a reliable proxy for gyrase binding sites, underscoring that stable binding and cleavage are mechanistically distinct processes. In turn, it highlights the need for more experiments to discern how DNA mechanics and gyrase binding affinity impact supercoiling propensity.

## DISCUSSION

### The mechanical code of DNA impacts DNA:protein interactions

Modern molecular biology has been driven by our understanding of how DNA sequence encodes information. While the genetic code encodes protein coding information, regulatory information is encoded in sequence *via*, for example, recognition sites for transcription factors, epigenetic modifications of DNA bases (promoter CpG methylation silencing transcription for instance), or the global 3D structure of the genome^1^. In this study, we have directly highlighted another route *via* which sequence can encode regulatory information controlling the formation of a large nucleoprotein complex – by controlling the mesoscale mechano-structural properties of the DNA polymer *via* the mechanical code.

The formation, remodelling, and dissolution of large nucleoprotein complexes involving extensively deformed DNA configurations is a problem that cells need to repeatedly solve in diverse contexts. For instance, architectural transcription factors like bacterial histone-like protein HU and Integration Host Factor (IHF) significantly bend DNA^33^, Structure Maintenance of Chromatin (SMC) proteins like cohesion extrude tight loops in the path of DNA^34^, nucleosomes^2^ and their remodelling rely on extensive DNA bending and wrapping, viral packaging of DNA involves tight bending^3^, and DNA repair factors like MutS have been purported to detect damage sites by assessing for local DNA flexibility via extensive DNA bending^1,35^.

Here we directly show that variations in DNA mechanics as driven by sequence variations that are (i) actually present within the genome, (ii) selected for on the basis of *in vitro* evolution, (iii) totally random, or (iv) synthetically created to have specific patterns, can all directly impact the interactions between DNA and gyrase – a large DNA-binding protein complex critically needed for all bacterial life. Coupled with our earlier more correlative exploration of how DNA mechanics impacts nucleosome organization^10,11^, our works likely indicate a broad role of the mechanical code in controlling diverse classes of critical DNA:protein interactions in diverse organisms.

### Novel roles of structural intermediates of the gyrase:DNA complex

Gyrase is the only known type II topoisomerase that can negatively supercoil DNA. This attribute is largely owing to the specialized CTDs that chirally wrap DNA, creating a positive loop in the path of DNA. Two negative supercoils are introduced by strand passage, which converts the positive loop into a negative loop. Therefore, historically, the role of the CTD was largely considered to aid chiral wrapping only. Our earlier single-molecule studies identified a new configurations of the nucleoprotein complex – Ω – which is long-lived, precedes formation of the chiral wrap, and involves extended DNA:CTD interactions. Exit from Ω was shown to be rate-limiting and involve a rearrangement of the enzyme-bound DNA in the shape of a chiral loop^21,22^.

What is the functional role of such a long-lived Ω state? Based on our current work, it is likely that CTD:DNA interactions are essential for stable binding of gyrase to DNA, because in absence of this, it is unlikely that extended regions of high intrinsic cyclizability would have been enriched by SELEX. This provides direct confirmation of the hypothesis that the enzyme dwells longest in a state involving extensive DNA:CTD interactions like Ω. It suggests that the long-lived Ω state, tuned by the mechanics of DNA itself, allows the enzyme to stably remain anchored to DNA, likely impacting processivity.

### Functional asymmetry within symmetric proteins and their substrates

Our observations support a hypothesis that the ability of one CTD tightly binding to DNA can compensate for weaker binding of the other. What advantage might there be to permit this? The gyrase mechanochemical cycle involves significant remodelling of the nucleoprotein complex, involving on-pathway transient release of wrapped DNA every cycle^21,30^, large rearrangements of wrapped DNA in Ω to create the looped shape, dynamic opening and closing of three protein:protein interfaces that act as gates for passing DNA strands, significant movements of the CTDs and their associated wrapped DNA as revealed by smFRET experiments^36^; and recently suggested extensive movements of the ATPase domains of the enzyme likely gated or times by conformational changes in the DNA^20^. It is therefore likely an evolutionary advantage for one CTD to be stably attached whereas the other to allow enough conformational flexibility of the wrapped DNA to permit necessary structural rearrangements during the cycle. Future single molecule or structural studies may be able to dissect the impact of DNA sequence on structural dynamics and mechanochemical coupling in gyrase.

Our work also adds to an evolving picture of symmetric proteins binding or interacting asymmetrically with their substrates. Nucleosomes are formed via the tight wrapping of 147 bp of DNA around the histone octamer. The octamer is highly symmetric, yet (i) unwrapping of DNA under disruption such as mechanical tension is highly asymmetric, mirroring the asymmetry in DNA flexibility between the two halves of the octamer, and (ii) asymmetric unwrapping of DNA from one half of the octamer stabilizes the wrap around the other half^37^. More recently, we also showed that promoter-proximal nucleosomes are usually formed on DNA substrates with asymmetric flexibility on either side of the known centre of the 147 bp nucleosomal binding region, and that for highly expressed genes, DNA on the promoter-proximal half is more flexible^11^. We hypothesized a similar functional role of permitting asymmetric binding – balancing the need for nucleosome positioning with disruption associated with transcription.

Apart from gyrase and nucleosomes, several other examples of functional asymmetries within symmetric proteins also exist. Symmetric dimers of transcription factors are known to bind with asymmetric DNA contacts^38^, symmetric AAA+ motor proteins undergo sequential asymmetric hydrolysis for directionality^39^, DNA repair or recombination factors often comprise symmetric filaments or dimers but display asymmetric DNA contacts of strand exchange dynamics^40^. The emergence of functional asymmetry within symmetric molecular assemblies is a recurrent theme in biology that may serve to enable directionality, regulation, and adaptability.

### Substrate DNA properties may impact structure and structural dynamics

Our observations suggest that substrate DNA can significantly impact enzyme binding, and likely structural dynamics as well, via its sequence-dependent mechano-structural properties. Earlier single-molecule characterizations of gyrase lead by us and others have heavily relied on using the MuSGS sequence. Here we show that MuSGS is one of the strongest gyrase binders that likely provides very high stability to DNA:CTD interactions, which in turn is fundamental to the overall catalytic activity of the enayme. In cell, however, gyrase interacts with DNA with diverse and distinct mechanical properties, such as those seen around mapped cleavage sites. Therefore, to what extent the characterized properties of gyrase like the structural dynamics of DNA wrapping in Ω and in the chiral wrapped state, the velocity and processivity of the enzyme, and its ability to work against existing superhelical tension, vary with the diversity of substrate DNA mechanics now needs to be investigated.

Even beyind gyrase, substrate DNA has largely been considered passive and lacking regulatory role in structural and biochemical characterizations of nucleoprotein complexes. This has historically led to entire fields being driven by experiments involving just a few different DNA sequences. Examples include the 601 strong nucleosome positioning sequence^41^ and the *Xenopus borealis* 5S rDNA nucleosome positioning sequence^42^ driving vast amounts of research on nucleosomes, the Mu phage Strong Gyrase Site being repeatedly used in characterizations of DNA gyrase^21,22^, the lacUV5 promoter being used as a standard promoter for RNA polymerase (RNAP) kinetic, footprinting, and structural studies^43,44^, the Integrative Host Factor (IHF) binding site in the phase λ attP region^44^ used as the canonical DNA sequence for studying DNA bending by IHF and the bacterial histone-like protein HU, and the human EMX1 locus as the canonical target for Cas9 assays in human cells^1,45^. This work highlights the critical need for broader exploration of sequence space in order to properly characterize how DNA-manipulating and DNA-interacting factors work. The methods developed here – SELEX and DNA-compete coupled with high-throughput predictions of DNA mechanics – lays a foundation for future large-scale explorations.

## ACKNOWLEDGEMENTS

We would like to thank Jonathan Heddle and Olivia Gittins for sharing expertise related to gyrase purification, Jonghan Park for help with applying the neural network models, and Zev Bryant for initial discussions. This work was funded by The Royal Society grant URF\R1\211659 to Aakash Basu-Biswas (also known as Aakash Basu), which also partly supported Bailey G. Forbes. Aakash Basu-Biswas (also known as Aakash Basu) is a Royal Society University Research Fellow. Rosella Pinckney was supported by the UKRI NLD Doctoral Training Programme. Aditi Biswas was supported by a Durham Doctoral Scholarship from Durham University. Heather Potter was supported by a Department of Biosciences Summer Studentship from Durham University. Sisi Liu was supported by funding from the China Scholarship Council.

## AUTHOR CONTRIBUTIONS

Bailey G. Forbes (BGF) and Aakash Basu-Biswas (ABB, also known as Aakash Basu) designed the experiments. BGF performed all experiments. BGF and ABB analysed the data and wrote the paper. Sisi Liu helped establish gyrase activity assays. Rosella Pinckney and Elizabeth R. Morris (ERM) purified active gyrase enzymes. Heather Potter helped with initial gyrase binding assay development. ABB conceived and supervised the project. ERM supervised gyrase purification and contributed to overall project development. All authors critically read the manuscript.

## USE OF GENERATIVE AI

Use of generative AI has been limited to assistance with writing custom python scripts for data analysis. All scripts generated using GenAI were checked for accuracy.

## Notes

### Competing Interest Statement

The authors have declared no competing interest.

