## Supplementary Notes, Tables, and Figures for "The mechanical code of DNA impacts its interaction with DNA gyrase"

[3]Also known as Aakash Basu

[*]Correspondence should be addressed to Aakash Basu-Biswas, also known as Aakash Basu (lead contact:) and Elizabeth R. Morris (co-corresponding author:).

**Supplementary note 2a: plotting details for Figure 2a**

The sequences of pSC101, the Mu phage genome, and PBR322, were obtained. The reported gyrase binding sites were present between coordinates 4580 – 4864, 17,765 – 18050, and 846 – 1126. The DNA sequence from 25 bp upstream of this window to 25 bp downstream was extracted and split into overlapping 50 bp fragments, each offset from the previous by 1 bp. Flexibilities of each of these fragments were obtained via the neural nets model[1]. Plotted are these flexibilities. For any 50 bp fragment that extends from position n to n+49 along the genome or the plasmid sequence, its mechanical obtained intrinsic cyclizability was assigned as the intrinsic cyclizability of DNA at position n+25 along the genome or plasmid sequence.

**Supplementary note 2c: plotting details for Fig. 2c**

For every round of SELEX, 10,000 unique reads were selected at random. These reads were all 167 bp long, comprising the left and right adapters (see SELEX methods) flanking the central variable 133 bp regions. For every round, each 167 bp sequence was split into 118 50 bp DNA fragments offset from each other by 1 bp. Intrinsic cyclizability values of each of these fragments was obtained using the neural nets based model. The predicted values were organized into a matrix of 10,000 × 118, where each row corresponds to one DNA sequence and each column to one 50 bp fragment position along that sequence. For each SELEX round, the column-wise mean of this matrix was calculated and plotted to generate the average flexibility profile. The x-axis positions for each of the 118 columns were assigned numerical values beginning at 25, increasing by 1 bp per column. In other words, for any 167 bp DNA sequence, the first 50 bp fragment (positions 1–50) contributes its predicted flexibility value at position 25, the second fragment (positions 2–51) at position 26, and so forth.

**Supplementary note 2d: plotting details for Fig. 2d**

To analyze the symmetry of DNA flexibility profiles among gyrase-binding sequences, we used the 10,000 randomly selected reads from the 7th round of SELEX. Each read was 167 bp long, comprising the constant left and right adapters flanking a 133 bp variable region.

(1) Segmentation and flexibility prediction

Each 167 bp sequence was divided into 118 overlapping 50 bp fragments, offset by 1 bp relative to the previous fragment. The intrinsic cyclizability of each fragment was predicted using our neural-network–based model.

(2) Smoothing of flexibility profiles

For each sequence, the resulting vector of 118 intrinsic cyclizability values was smoothed using a Gaussian filter with a standard deviation $\sigma = 2$ bp to suppress noise while preserving the broad shape of the profile. Smoothing was performed using python scripts using scipy.ndimage.gaussian_filter1d from SciPy 1.10.0.

(3) Peak detection and classification

Peaks were identified in each smoothed profile using the python scripts that call the function scipy.signal.find_peaks from SciPy 1.10.0, with the height threshold set to the mean + one standard deviation of that profile. This algorithm can detect any number of peaks along the profile that satisfy the threshold—there was no constraint on the number of peaks per side.

Detected peak positions were then partitioned into those lying before the midpoint, and those after the midpoint, of the 167 bp DNA region. Sequences containing at least one left and one right peak were labeled *"both"*. Sequences with only left-side peaks were labeled *"left"*. Sequences with only right-side peaks were labeled *"right"*. Sequences with no detected peaks were labeled *"none"*.

(4) Averaging within each class

Profiles were grouped by these four categories. For each class, the mean and standard deviation of intrinsic cyclizability were computed at every position across the 118 columns, producing an average ± SD profile for that class. These 4 profiles for the 4 classes are plotted.

**Supplementary note 3b: categorical spectral analysis, plotting details for Fig. 3b, phasing**

<u>Categorical spectral analysis</u>

To analyze periodic patterns in the base composition of DNA sequences, we performed Fourier analysis. Taking the Fourier transform of a DNA sequence inherently requires some strategy for converting base composition into numeric data. The exact numeric representation (for example, A/T = 0 and G/C = 1) affects the calculated power spectral density[2].

We therefore performed categorical spectral analysis which treats individual sequences as a sequence of various 'categories' (A/T or G/T), and numerically represents categories in an unbiased manner. We performed categorical (Voss-type) spectral analysis separately for the AT and GC base categories for every given sequence. Every sequence is first converted into two indicator binary traces of the same length: an AT indicator trace, where positions corresponding to A or T are assigned a value of 1 and all others 0, and a GC indicator trace, where positions corresponding to G or C are assigned a value of 1 and all others 0. All analyses were performed in Python 3.10, using NumPy (v1.24.3), SciPy (v1.10.0), and Matplotlib (v3.7.1).

Each indicator trace was detrended by subtracting its mean value, ensuring that spectral power reflects oscillatory variation rather than base composition bias. To suppress edge artifacts, a Hann window was applied prior to the Fourier transform.

After detrending and applying a Hann window, the indicator traces for AT and GC bases were converted from the spatial domain to the frequency domain using the Fast Fourier Transform (FFT). This operation decomposes the base-composition signal into a sum of sinusoidal components of different spatial frequencies (cycles per base pair). The squared magnitudes of the resulting complex Fourier coefficients were used to compute the power spectrum for each sequence, reflecting the relative strength of periodic A/T or G/C patterns along the DNA.

For every sequence, the total categorical power spectrum was calculated as the sum of the individual A/T and G/C power spectra at each frequency. The total categorical power spectra from all sequences were averaged frequency-wise to obtain the quantity plotted as 'Power' in Fig. 3b.

<u>Left and right panels in fig. 3b:</u>

In the left panel, Power was obtained from the entire 133 bp DNA sequence, averaged over all 10,000 sequences in each round of SELEX. In the right panel, the same set of sequences was chosen (for round 7 SELEX only), but the sequences were split into left 60 bp (positions 1-60 along the 133 bp region) and right 60 bp (positions 74 -133 along rhe 133 bp region). This was done to investigate whether periodic oscillations are more pronounced on one half of the molecules or not.

<u>Phasing leading to apparent GC-content oscillations:</u>

In Fig. 3a, the plot of the mean GC content as a function of position, averaged across all 10,000 selected reads from among the reads of the $7^{th}$ round of SELEX visually show more pronounced periodic oscillations on the "left" half of the molecule. When considering a large set of individual periodic sequences, all of the same length, the per-position average of these sequences will also be periodic only if individual sequences are phase matched. We therefore speculate that the adapter sequence on the left half of the molecule likely imposes a specific phase because it might have a strongly preferred binding site along the CTD. The same may not be true for the right adapter.

Our power spectral analysis method, however, takes the Fourier transform of individual sequences, and averages the individual power spectra to obtain the final averages power as a function of peiordicity plot. Therefore, a peak indicates that individual sequences, on average, have a strong periodic signature at that specific period. This is not equivalent to taking an average sequence first, and then taking a fourier transform of the average sequence. Mathematically, the act of averaging, and taking fourier transform, do not commute, but this has been the cause of much confusion in the literature, especially in relation of periodic sequence features along known nucleosome binding sites[2].

**Supplementary Note A: identification of SELEX-enriched 6-mers**

To identify sequence motifs enriched during SELEX, we compared the frequencies of all possible 6-mers between 10,000 randomly selected sequences from round 7 and 10,000 sequences from round 0. All analyses were performed in Python 3.10 using NumPy (1.24.3), Pandas (2.0.1), and SciPy (1.10.0). For each DNA sequence, all valid 6-mers were enumerated using a sliding window advanced by one base at a time. This produced complete non-overlapping k-mer count dictionaries for the two datasets (the 10,000 randomly selected unique reads in round 7 and round 0 of SELEX). Reverse complements were not merged, so enrichment was computed for each 6-mer orientation separately.

For every possible 6-mer, we built a 2x2 table. Row 1 stores the number of times that 6-mer occurs in the roung 7 pool and in the round 1 pool. Row 2 stores the number of other 6-mers that occur in these pools (i.e. the number of times the 6-mer did not occur). Fisher's exact test (scipy.stats.fisher_exact, option alternative='greater') was used to then evaluated whether the proportion of that 6-mer among all k-mers was significantly higher in the round-7 pool than in the round-0 pool; the resulting one-sided *p*-value quantifies the likelihood of observing such enrichment by chance.

Because thousands of different 6-mers were tested, we corrected the resulting *p*-values for multiple comparisons using the Benjamini–Hochberg false discovery rate (FDR) method. This adjustment controls the expected proportion of false positives among all k-mers identified as significantly enriched. The FDR-adjusted *p*-values (referred to as *q*-values) were calculated using a custom implementation of the standard Benjamini–Hochberg ranking procedure.

The resulting statistics for every k-mer (counts in both pools, frequencies, odds ratio, p and q values, and $\log_2$enrichment were compiled into a spreadsheet and sorted by decreasing log2enrichment. The top 20 most significantly enriched 6-mers were used for subsequent analysis).

**Supplementary Table 1: list of SELEX-enriched 6-mers**

Identified 6-mers, their $\log_2$ enrichment values, p, and q, based on the method discuss above in Supplementary Note A is presented below:

| 6-mer | log2 enrichment | p | q |
|---|---|---|---|
| **GCTCCT** | 2.0 | 7E-76 | 1E-73 |
| **CTCCTA** | 1.9 | 2E-52 | 2E-50 |
| **GGCGTT** | 1.7 | 1E-142 | 1E-139 |
| **TAGGGC** | 1.6 | 9E-119 | 6E-116 |
| **GGCTCC** | 1.6 | 2E-55 | 2E-53 |
| **GGCTCT** | 1.6 | 3E-76 | 8E-74 |
| **GGGGCT** | 1.6 | 1E-165 | 2E-162 |
| **GGCGCT** | 1.6 | 2E-76 | 6E-74 |
| **TCCTAG** | 1.6 | 3E-44 | 2E-42 |
| **GAGCCT** | 1.5 | 3E-53 | 3E-51 |
| **GGCCCT** | 1.5 | 3E-47 | 3E-45 |
| **GGGGGG** | 1.5 | 1E-258 | 4E-255 |
| **GCGTTT** | 1.5 | 1E-90 | 4E-88 |
| **GGGCGT** | 1.5 | 8E-125 | 7E-122 |
| **GCCCCT** | 1.5 | 1E-23 | 4E-22 |

| | | | |
|---|---|---|---|
| **CGCTGT** | 1.4 | 9E-51 | 8E-49 |
| **TGGGGC** | 1.4 | 5E-116 | 2E-113 |
| **AGGGCG** | 1.4 | 2E-84 | 7E-82 |
| **GCGGCT** | 1.4 | 1E-59 | 2E-57 |
| **GCTGTG** | 1.3 | 2E-76 | 5E-74 |

Because the motif coverage of the above motifs was significant only in the two regions of CTD binding flanking the centres of the DNA fragment, we attempted to re-obtain enriched 6-mers, this time focusing only on this region for comparing between 6-mers in round 7 (final round) vs round 0 (positions 15 –35 along the 133 bp central variable region of the 167 bp molecules). The resulting identified 6-mers, along with their $\log_2$ enrichment values, p, and q, are as below, and indicate higher enrichment values:

| **6-mer** | **log2 enrichment** | **p** | **q** |
|---|---|---|---|
| **GCTCCT** | 4.2 | 2E-59 | 2E-56 |
| **CTCCTA** | 3.5 | 6E-42 | 4E-39 |
| **CCCCTT** | 3.3 | 1E-19 | 2E-17 |
| **GGCTCC** | 3.0 | 4E-36 | 2E-33 |
| **TCCTAG** | 2.9 | 7E-34 | 2E-31 |
| **CCCCCT** | 2.9 | 7E-13 | 4E-11 |
| **CTAGGG** | 2.8 | 3E-62 | 7E-59 |
| **TATCCC** | 2.8 | 9E-11 | 3E-09 |
| **ATCCCC** | 2.7 | 2E-11 | 9E-10 |
| **GCCCTA** | 2.7 | 1E-19 | 2E-17 |
| **TTCCTA** | 2.7 | 2E-19 | 3E-17 |
| **GCTCTG** | 2.7 | 1E-33 | 5E-31 |
| **CCTAGA** | 2.6 | 2E-22 | 3E-20 |
| **GGCCCC** | 2.6 | 8E-15 | 5E-13 |
| **CTTTGC** | 2.6 | 2E-23 | 3E-21 |
| **CCTATT** | 2.6 | 1E-20 | 1E-18 |
| **GCCCCC** | 2.6 | 5E-09 | 1E-07 |
| **TCCTAT** | 2.6 | 2E-21 | 2E-19 |
| **GATCCC** | 2.6 | 4E-17 | 3E-15 |
| **AGCCCT** | 2.5 | 3E-17 | 2E-15 |

For all subsequent analysis, only the first table of enriched 6-mers (that were identified from the full 133 bp regions) was considered.

**Supplementary note 3d: plotting details for Fig. 3d**

GC-matched random sequences were generated from the 10,000 selected DNA sequences from round 7 of SELEX. For every sequence, we replaced Gs or Cs randomly with Gs or Cs, likewise for As and Ts. Thus, every generated sequence matches the G/C distribution of the SELEX sequence used to generate it, but was otherwise random. These generates sequence were again flanked by the same 17 nt left and right adapters. Mean intrinsic cyclizability along these sequences was plotted, as done in Fig. 2c.

Motif-matched random sequences were similarly generated from the same 10,000 sequence set. For every sequence, at each of the 133 positions between the left and right adapters, a base was marked as 'covered' if it was part of any of the identified 20 6-mer motifs. Then, a random sequence was generated which still retained the same coverage profile. Mean intrinsic cyclizability was plotted as in Fig. 2c.

For the cleavage sequences, all previously reported 4,627 Cfx-induced gyrase cleavage sites along E. *coli*[3] were considered, and the 133 bp DNA sequence flanking these sites were extracted. These 133 bp sequences were then flanked by the same 17 nt left and right adapters to generate 167 bp sequences. Mean intrinsic cyclizability as a function of position was plotted for these sequences, exactly as done in Fig. 2c.

**Supplementary Figure S1:**

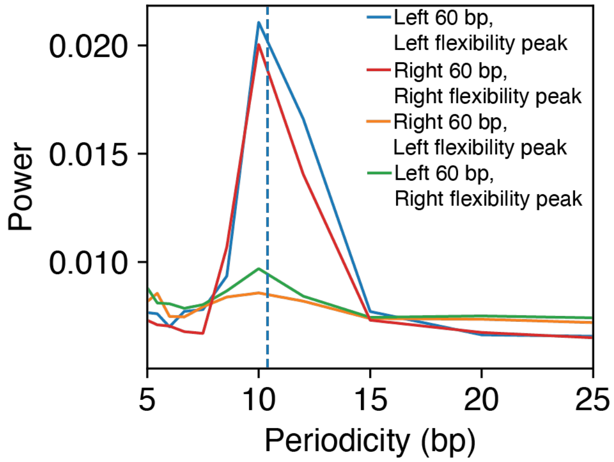

***Fig. S1:*** *The 10,000 randomly chosen unique reads from round 7 of SELEX were split into the 1 categoris depending on whether the sequences had a peak in intrinsic cyclizability on the left half, on the right half (Fig. 2d). Sequences with flexibility peaks on both halves, or those with no detected peaks, were left out from this analysis. For each category individually, we calculated and plotted the mean categorical power for GC content oscillations, when considering only the left 60 bp of the sequences, or the right 60 bp of the sequences. We find that sequences grouped to have a flexibility peak on the "left", also have a discernible peak in power on the left half only, and likewise for sequences classified as having a flexibility peak on the "right". Therefore, asymmetries in DNA flexibility mimic asymmetries in the extent of periodic GC content oscillations.*

**Supplementary Figure S2: in silico evolution to generate flexibility matched random sequences**

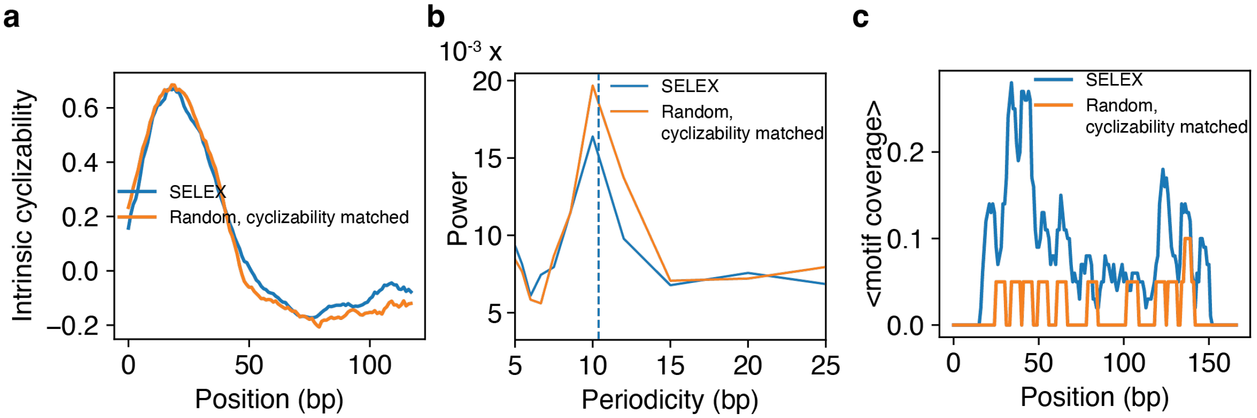

<u>*In silico* evolution to generate DNA sequences with the specific intrinsic cyclizability profile along its length.</u>

To generate a DNA sequence whose intrinsic cyclizability profiles matches some given profile, we start with a totally random DNA sequence. We consider all possible single point mutation variants of this sequence, calculate the profiles of each of these variant sequences (by splitting the sequence into overlapping 50-mers and using the neural nets model to predict their intrinsic cyclizabilities), and select the one with the least RMS deviation from the profile we are trying to match. We then consider all possible single point mutations of this selected sequence and repeat the process. This iteration, comprising single point mutation and selection, was carried out *in silico* 15 times.

<u>Panel (a):</u>

Plotted in panel (a) is the mean intrinsic cyclizability as a function of position along these generated 20 flexibility-matched DNA sequences, and along the original 100 SELEX sequences whose flexibility profiles we were trying to match. Close match shows we were successful in generating otherwise random DNA sequences that matched a certain intrinsic cyclizability profile.

<u>Panel (b):</u>

Using categorical spectral analysis, we calculated the total Fourier power as a function of periodicity, associated with oscillations in GC content along the 100 SELEX sequences and the 20 randomly generated sequences. This was done exactly as in Fig. 3b. Only the first 60 bp was considered, as the flexibility peak was on the left side of the molecule, and we have already demonstrated that flexibility asymmetry mimics GC periodicity asymmetry (Supplementary Fig. S1). Dashed line represents 10.4 bp. We find comparable ~10 bp periodic signaturs among both classes of sequences, suggesting that GC content oscillations is an automatic consequence of selection for intrinsic cyclizability.

<u>Panel (c):</u>

Mean motif coverage profiles (as in Fig. 3c) of the 100 SELEX sequences, and the 20 cyclizability-matchd random DNA sequences, along the central 133 bp region. Selection for DNA cyclizability does not ensure the same coverage density of the identified 6-mer motifs.

**Supplementary Figure S3:**

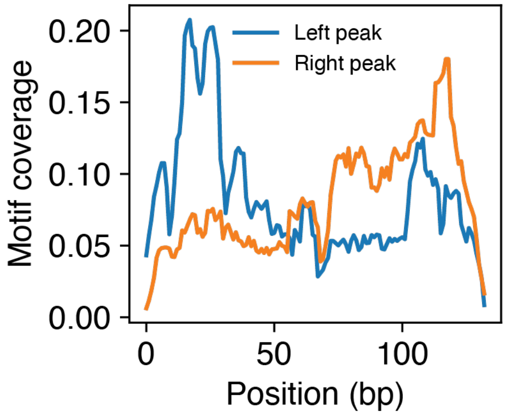

We determined whether asymmetries in DNA cyclizabilities are correlated with asymmetries in the coverage densities of the identified 6-mer motifs. We considered the 10,000 randomly selected unique reads from the 7$^{th}$ round of SELEX and considered those classified as having cyclizability peaks on the left or the right halves only. We obtain the motif coverage density averaged across these groups separately (as done in fig. 3c). We find that sequences classified as having cyclizability peaks on the left half also have higher motif coverage on the left half alone, and vice versa.

**Supplementary note 4c: plotting details for Fig. 4c**

As described, 400 DNA sequences were artificially synthesized, with 20 sequences in each of 20 groups. DNA sequences within a group were created to have a certain intrinsic cyclizability profile along their lengths. The mean intrinsic cyclizability as a function of position, averaged over the 20 sequences in each group, is shown in Fig. 4a.

Left panel of Fig. 4c: Each point represents DNA sequences within a group. The y coordinate is the mean (error bars are s.e.m.) of the binding scores of the 20 sequences within that group. For each sequence in the group, its intrinsic cyclizability profile (intrinsic cyclizability as a function of position along the 167 bp region) is obtained as described in Fig. 2c, and the average cyclizability between positions 40 – 50 bp from the left edge is calculated. This is a measure of the "height" of the single intrinsic cyclizability peak on the left half of the molecule. The average of these average cyclizability values across all 20 sequences of the group is calculated and plotted as the x-coordinate of every point.

Right panel: Here, for DNA sequence belonging to groups 11-20, the x-coordinate is the sum of "heights" of both the peaks, spannings locations 40 – 50 bp (for the "left" peak) and 115 – 125 bp (for the "right" peak) along the 167 bp region.

**Supplementary note B: linear regression model to predict biding score**

We modelled the binding score (B) of each 133 bp sequence as a linear combination of its maximum cyclizability along the 167 bp region (when the 133 bp region is flanked by the constant adapters) and overall GC content along the central 133 bp variable region:

$B = \beta_0 + \beta_1 \times$ (max cyclizability) $+ \beta_2 \times$ (GC content) ------ (1) (see supplementary note B)

Cyclizability is defined only for 50 bp fragments. For any 133 bp DNA sequence, we flanked it by the 17 bp left and right adapters (as done in SELEX), and tiled the resulting 167 bo segment into 167-50+1 = 118 overlapping 50 bp chunks, with each chunk offset from the previous by 1 bp. The maximum flexibility of the original 133 bp sequence is simply the maximum of these 118 values. The GC content is the total number of Gs and Cs in the 133 bp segment. Fitting this model to the 400 random sequences in the library produced coefficients that quantitatively captured the observed correlations. The fitted coefficients were $\beta_0 = -0.30 \pm 0.08$, $\beta_1 = +0.28 \pm 0.03$, and $\beta_2 = -0.011 \pm 0.001$ (mean ± s.e.m.). Both predictors were highly significant ($P < 10^{-9}$). As expected, $\beta_1$ was positive and $\beta_2$ was negative, confirming that increased flexibility enhances binding whereas higher GC content suppresses it.

Once fitted, the model was applied to all sequences in the library to compute predicted binding scores using their corresponding values of max cyclizability and *GC* content.

Predicted values were then compared with the experimentally measured binding scores across all sequence categories, including random sequences (which were used to train the model), synthetic flexibility-modulated sequences (groups 1-20), cleavage-site–derived sequences, and SELEX-enriched sequences. The model accurately reproduced the trend in observed dependence of binding on flexibility and GC content across all groups (Fig. 5c). However, sequences enriched through SELEX consistently showed higher binding scores than predicted, suggesting that additional sequence features—such as specific GC-rich motifs as identified—enhance gyrase binding beyond what is captured by the model.

**Supplementary note 5g: plotting details for Fig. 5g**

The 1,797 2,001 bp DNA sequences flanking the reported 1,797 Transcription Start Sites (TSSs) in *E. coli*[4] from 1,000 bp upstream to 1,000 bp downstream were extracted from the genome sequence of *E. coli* and from the list of TSS locations (also listed in RegulonDB (https://regulondb.ccg.unam.mx/)). Strand information of the TSSs was taken into account in determining upstream or downstream direction. Each 2,001 bp DNA sequences was parsed into 1,835 overlapping 167 bp segments, each segment offset from the previous by 1 bp. For each of these segments, we first independently obtained the number of Gs and Cs along the central 133 bp region of the 167 bp fragment it as the GC content of that segment. We then parsed this 167 bp fragment into 118 overlapping 50 bp chunks, each offset by 1 bp from the previous, and used our neural nets model to predict intrinsic cyclizability of all these chunks. We obtained the maximum value of these 118 predictions and noted it as the maximum cyclizability of the chunk. We then used our model (equation 1 of the main text) to predict intrinsic cyclizability of the 118 bp chunk, based on the GC content and maximum flexibility values as inputs.

For every 2,001 bp sequence spanning a TSS, the 1,835 predicted binding scores are arranges as numbers along the rows of a matrix. Plotted in Fig. 5g is the column-wise average of this matrix. The x coordinate refers to the centres of the 167 bp fragments along the original 2,001 bp region extending from -1,000 to +1,000. As an example, since the first 167 bp chunk of any 2,001 bp fragment extended from position -1,000 to -834, its predicted binding score was assigned the x value (-1,000+floor(167/2)) bp = (-1,000+83) bp = -917 bp.
